# The Temporal Dynamics of Opportunity Costs: A Normative Account of Cognitive Fatigue and Boredom

**DOI:** 10.1101/2020.09.08.287276

**Authors:** Mayank Agrawal, Marcelo G. Mattar, Jonathan D. Cohen, Nathaniel D. Daw

## Abstract

Cognitive fatigue and boredom are two phenomenological states that reflect overt task disengagement. In this paper, we present a rational analysis of the temporal structure of controlled behavior, which provides a formal account of these phenomena. We suggest that in controlling behavior, the brain faces competing behavioral and computational imperatives, and must balance them by tracking their opportunity costs over time. We use this analysis to flesh out previous suggestions that feelings associated with subjective effort, like cognitive fatigue and boredom, are the phenomenological counterparts of these opportunity cost measures, instead of reflecting the depletion of resources as has often been assumed. Specifically, we propose that both fatigue and boredom reflect the competing value of particular options that require foregoing immediate reward but can improve future performance: Fatigue reflects the value of offline computation (internal to the organism) to improve future decisions, while boredom signals the value of exploration (external in the world). We demonstrate that these accounts provide a mechanistically explicit and parsimonious account for a wide array of findings related to cognitive control, integrating and reimagining them under a single, formally rigorous framework.

## Introduction

Learning is one of the most widely studied processes in all of cognitive psychology. New tasks are often difficult, but they become easier – and subsequently, we perform them better – with practice (Posner & Snyder, 1975; Schneider & Shiffrin, 1977; Shiffrin & Schneider, 1977; Anderson, 1987). Computational models of learning propose that this is the result of minimizing prediction errors, and can be captured by connectionist models using backpropagation and/or reinforcement learning models using temporal difference learning (Sutton, Barto, et al., 1998; J. D. Cohen, Dunbar, & McClelland, 1990; Daw, Niv, & Dayan, 2005).

However, to date, these models and most other formal theories of learning have largely failed to address the ubiquitously recognized subjective states of *cognitive fatigue* and *boredom*, and the changes in objective performance associated with these. Most theories predict that, with practice, there should be monotonic improvements in performance. In accord with this prediction, greater practice does generally lead to progressive improvements in performance. For example, a participant training on a task for an hour every day will usually perform better after two weeks. Yet, most learning models do not take account of how the temporal characteristics of practice influence performance. They would naively predict that performance at the end of fourteen straight hours of practice is comparable to that at the end of the same amount of practice carried out periodically over two weeks^1^. This is unlikely to be true (Arai, 1912; Huxtable, White, & McCartor, 1946; N. H. Mackworth, 1948; Van der Linden, Frese, & Meijman, 2003; Healy, Kole, Buck-Gengler, & Bourne, 2004; Lorist, Boksem, & Ridderinkhof, 2005; Warm, Parasuraman, & Matthews, 2008; Haager, Kuhbandner, & Pekrun, 2018). Specifically, after prolonged task engagement, it is all but certain that participants will feel *fatigued* or *bored*, and make considerably more errors (if they continue to perform for the full duration at all).

Fatigue has often been attributed to the consumption, and consequent diminution, of some resource (e.g., metabolic; Baumeister, Bratslavsky, Muraven, & Tice, 1998; Baumeister & Vohs, 2007), by analogy to the case of physical fatigue. In contrast, Kurzban, Duckworth, Kable, and Myers (2013) proposed that exerting cognitive control (limitations of which may potentially characterize fatigue and boredom) may instead be accompanied with a sensation that signals opportunity costs that are a consequence of the limitation in the number of tasks that can be performed concurrently. That is, with the passing of time, it becomes increasingly possible that behaviors other than the one currently being performed offer opportunities for greater reward, and to which it would be more valuable to switch behavior (Bench & Lench, 2013; R. Hockey, 2013; Inzlicht, Schmeichel, & Macrae, 2014). However, this broad approach admits of many specific models. In addition to focusing on particular sensations (e.g., boredom), a fully specified opportunity cost model must satisfy two criteria: (1) it must define the nature of the alternative behaviors that give rise to the opportunity cost(s); and (2) it must account for the temporal dynamics of the phenomena it is meant to explain (*e.g.*, the increase in fatigue and boredom over time). The goal of the present research is to specify such a formal theory for the cases of fatigue and boredom.

### Overview

Here, we propose that one important class of opportunity costs arises from an intertemporal choice every agent must make: whether to sacrifice current reward in order to gather information that will result in greater reward later. Information gathering has value, which (if foregone, to instead pursue proximate reward) imposes an opportunity cost. As stated, this is the classic explore-exploit dilemma. However, importantly, we extend this analysis to consider two different types of information gathering actions. One corresponds to the standard treatment of explore-exploit: seeking out new opportunities in the external world, which can improve later choices. The second reflects an internal counterpart: offline processing by which one learns by thinking and mental simulation, again to compute decision variables that can improve future decisions. Both reflect a similar type of tradeoff between on-task performance and information gathering, but their values (and conversely, the opportunity costs for not pursuing them) have different dynamics with training. We identify the subjective feelings of boredom and fatigue with these specific, quantifiable decision variables which we propose our brain must compute and use for optimally allocating control.

## Roadmap

Part 1 proposes that fatigue adaptively signals the value of rest, and that the value of rest is derived from offline (internal) processing mechanisms such as hippocampal replay. We demonstrate how casting replay (a covert computational operation) as an instrumental action competing with overt behaviors leads to nontrivial dynamics of arbitration between replay and physical action in order to maximize future reward. Part 2 proposes that boredom tracks the value of exploring, here playing out via competition between different classes of overt action: information-seeking vs. exploitative. We offer a model that enables agents to navigate the explore-exploit tradeoff, expose its analogy to the case of replay, and demonstrate how the temporal dynamics of uncertainty lead agents to oscillate between different tasks. Finally, Part 3 integrates the two mechanisms of replay and exploration and examines new insights and problems arising from their interaction. Once viewed in the same framework, the internal and external actions so far discussed separately can also interact, with consequences for their value, such as in the case of planning to explore. We examine some of these cases and speculate whether they may relate to additional subjective phenomena such as mind-wandering. In summary we propose that, rather than reflecting hindrances as is often assumed, fatigue and boredom reflect control optimizations that track the values of replay and exploration, respectively, and are used by agents to maximize long-term reward. A schematic of this decomposition is illustrated in Figure 1.

**Figure 1.**
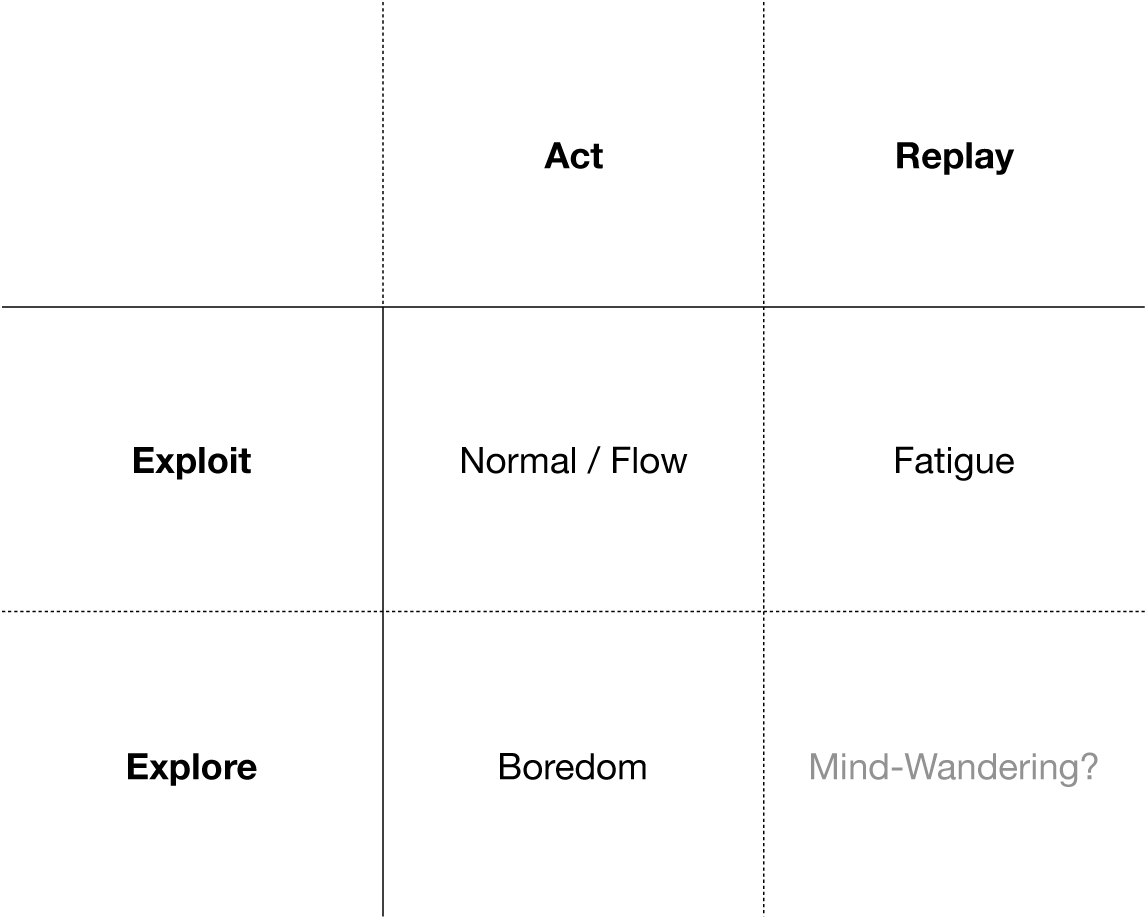
This paper partitions the constraints associated with the duration of cognitive control into two dimensions: action vs. replay and exploration vs. exploitation. Part 1 investigates fatigue, which we propose to be a signal for the value of replay while Part 2 investigates boredom, which we propose to be a bias towards exploration. Part 3 integrates the mechanisms of replay and exploration; whether this corresponds to a currently studied phenomenology is unknown and is an important direction for future research.

Before we formally introduce our models of fatigue and boredom, we present background on cognitive control and reinforcement learning.

## Background

### Rational Models of Cognitive Control

Cognitive fatigue has long been studied in psychology (E. Thorndike, 1900; Dodge, 1917), with one influential account, ‘ego depletion,’ suggesting that it reflects the consumption and subsequent diminution of a metabolic resource (Baumeister et al., 1998; Baumeister & Vohs, 2007). Recently, however, this hypothesis has been called into question (Kurzban et al., 2013); and multiple meta-analyses have challenged its empirical foundation, suggesting that the associated studies overestimated null effects (Carter, Kofler, Forster, & McCullough, 2015; Hagger et al., 2016; Randles, Harlow, & Inzlicht, 2017). But beyond problems with the particular experiments to which it was applied, the metabolic hypothesis itself also seems mechanistically flawed: what exactly is this metabolic resource? Glucose has often been proposed, but there does not seem to be a relationship between executive function and glucose levels (Messier, 2004; Raichle & Mintun, 2006; Gibson, 2007; Molden et al., 2012; Schimmack, 2012). In fact, some of the largest glucose demands in the brain arise from visual processing (Newberg et al., 2005), leading this account to predict, for instance, that face recognition should be more fatiguing than multi-digit arithmetic, though the opposite seems true in everyday life.

In contrast, normative models of cognitive control (Shenhav, Botvinick, & Cohen, 2013; Kurzban et al., 2013; Lieder & Griffiths, 2020) propose that performance variation arises from rational balancing of the costs vs. benefits of different control strategies, rather than biological resource limitations. Although these accounts vary as to how they operationalize control (and thus what ultimately makes it costly or limited), the cost-benefit framing implies that performance decrements due to fatigue or boredom can be countered with incentives, shifting the tradeoff.

The current theory instantiates a rational model to explain fatigue and boredom. Agents’ actions in our model span two dimensions: physical vs. mental and exploratory vs. exploitative (Figure 1). Because an agent can only perform one action at a time (and in particular, because the mental actions we consider are assumed to exclude physical ones, for reasons later justified), each action (including covert, internal ones) comes with the opportunity cost of foregoing all other actions. The goal of the rational controller is to identify the sequence of actions that maximizes reward.

### Reinforcement Learning

Reinforcement learning (Sutton et al., 1998; Daw et al., 2005) offers an integrative computational framework in which to implement a rational agent. Sequential decision problems in reinforcement learning settings are often modeled through a Markov decision process (MDP), a 5-tuple (*𝒮, 𝒜, ℛ, 𝒫, γ*), in which *𝒮* is the set of states, *𝒜* is the set of actions, *ℛ*(*s*) is the reward received in state *s*, *𝒫*(*s, a, s′*) is the probability of transitioning to from state *s* to state *s′* using action *a*, and *γ* is the discount factor. The policy *π* : *𝒮* ↦ *𝒜* determines with what probability an agent should perform action *a* when in state *s*.

#### Model-Free vs. Model-Based Learning

Two main classes of algorithms have emerged in reinforcement learning: model-free and model-based learning (Sutton et al., 1998; Daw et al., 2005). These are exemplified by two different approaches for using trial-and-error experience to estimate the value of candidate actions so as to guide choices toward better options. Formally, we consider the *value function*, the expected cumulative future discounted reward *Q*(*s, a*) for taking some action *a* in state *s*.

Model-free methods, such as Q-learning (Watkins & Dayan, 1992) estimate this function directly from experienced rewards over experienced state trajectories (Montague, Dayan, & Sejnowski, 1996; Schultz, Dayan, & Montague, 1997; Schultz, 1998). Here, an agent maintains an estimated function *Q*(*s, a*), and updates it after every experienced state-action-reward-state transition (*s, a, r, s′*), according to the temporal difference learning backup rule:

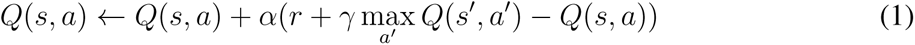

in which *α* refers to the learning rate of the agent. In contrast, model-based learning (Daw et al., 2005; Solway & Botvinick, 2012) learns an internal model of the environment (i.e. estimates the one-step dynamics *𝒫* and rewards *ℛ*). Such a model can be used to *compute Q*(*s, a*) by iterating steps and aggregating expected reward, formalizing a sort of mental simulation.

A main difference between these approaches is the computational work required to evaluate an action: model-based evaluation requires extensive internal iteration prior to action, whereas model-free values can be simply retrieved. Conversely, model-based evaluation is generally more accurate and flexible; this is because individual updates from Eq. 1 teach the agent about local rewards and costs, but working out their consequences for longer-run action-outcome relationships requires additional mental simulation (or many more experiential updates). Importantly, such mental simulation can “teach” the agent how to make better choices in future, but without actually collecting new information from the environment — instead, by discovering the consequences of information already known. For these reasons this computational distinction has been employed in neuroscience as a model of the tradeoffs between thinking and acting (Daw et al., 2005; Keramati, Dezfouli, & Piray, 2011). The general idea is that the brain can either act immediately according to (fast, potentially inaccurate) model-free values, or spend time computing better (more accurate) model-based ones. This leads to a speed-accuracy tradeoff and a rational account of many phenomena of habits, automaticity, compulsion, and slips of action (Daw et al., 2005; Keramati et al., 2011; Otto, Gershman, Markman, & Daw, 2013; Otto, Raio, Chiang, Phelps, & Daw, 2013; S. W. Lee, Shimojo, & O’Doherty, 2014; Keramati, Smittenaar, Dolan, & Dayan, 2016; Kool, Cushman, & Gershman, 2016; Kool, Gershman, & Cushman, 2017; Sezener, Dezfouli, & Keramati, 2019).

The current theory employs a finer grained version of this idea, based on the Dyna framework (Sutton, 1991), which learns values from experience using Eq. 1, but also makes decisions about whether to improve these using individual steps of model evaluation and, if so, which ones (Mattar & Daw, 2018). The core tradeoff — whether to act, or delay action to produce more accurate evaluations — and the logic of its cost-benefit resolution remain the same. Mattar and Daw (2018) proposed that the brain implements the steps of model evaluation by replaying trajectories in hippocampus. This theory will be the basis of our analysis in Part 1, where we suggest fatigue tracks the value of such replay.

#### Exploration vs. Exploitation

Reinforcement learning also offers an analysis of the explore-exploit tradeoff. When picking which restaurant to visit, whether to date a potential partner, or what research programs to pursue, humans must decide whether to *exploit* options they know are rewarding or forego those to *explore* new options that may potentially be even more rewarding. The explore-exploit tradeoff has long been studied in computer science and has recently attracted increasing attention in psychology and neuroscience (Kaelbling, Littman, & Moore, 1996; Sutton et al., 1998; Daw, O’doherty, Dayan, Seymour, & Dolan, 2006; J. D. Cohen, McClure, & Yu, 2007; R. C. Wilson, Geana, White, Ludvig, & Cohen, 2014; Mehlhorn et al., 2015; Gershman, 2018; Schulz et al., 2019).

The value of exploratory actions, in principle, is that the agent may learn something from them that improves their future choices — and thus their future earnings. The classic decision-theoretic analysis of the explore-exploit dilemma in problems such as bandit tasks (Gittins, 1979) attempts to quantify this long-run value directly by computing actions’ expected future value taking into account the possible effects of learning on later choices and rewards. This gives rise to a difference, for each action, between its nominal value *Q* based on current knowledge, vs. its expected long-run value including the improvement due to learning. For our purposes, we can generically express this as the sum of a baseline value *Q* and an additional increment, called value of information (VOI):

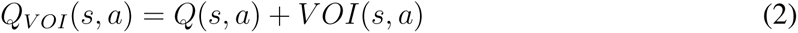

In practice, the VOI depends on the task: for instance, how it is broken up into repeated episodes, and what is shared between them (and/or other tasks) that can be learned to improve later performance. In general, although it can be defined formally, computing it exactly is typically intractable except for particular special cases. However, there are many heuristics and approximations to it. These can be added to nominal value *Q* to help the agent pursue exploratory behavior over immediate reward in circumstances in which that leads to greater overall (i.e., long-term) reward. A typical example is the Upper Confidence Bound (UCB) algorithm (Auer, Cesa-Bianchi, & Fischer, 2002), which proposes the VOI in a multi-armed bandit setting to be

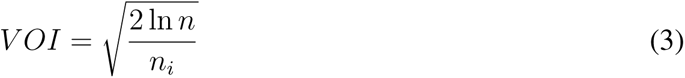

in which *n* is the total number of trials and *n_i_* is the number of trials arm *i* has been selected. According to this formula, *VOI* increases as time passes, and decreases with the number of times a given action is selected. Like many such heuristics, this quantity is a rough proxy for uncertainty about the value *Q*, which in turn measures how much can be learned about the task. This framework will be the basis of our analyses in Part 2, where we suggest increasing boredom reflects a bias towards exploration.

## Part 1: Cognitive Fatigue

A nearly ubiquitous observation is that, as we exert mental effort, we experience fatigue and eventually want to take a break. Furthermore, after taking a break, we may feel rejuvenated and willing to perform the task again. Fatigue has long been associated with rest (Kool & Botvinick, 2014; Müller & Apps, 2019), with Edward Thorndike defining fatigue as “that diminution in efficiency *which rest can cure* (emphasis ours)” over one hundred years ago (E. L. Thorndike, 1912). Here, we propose the value of rest is derived from offline computational processes such as hippocampal replay. But first, we summarize the evidence any rational theory of fatigue must explain.

### Empirical Findings

We identify three canonical effects in the literature a rational model of fatigue needs to explain: (1) why rest is valuable; (2) when an agent should switch between rest and action; and (3) why difficult tasks are more fatiguing than easier tasks.

#### 1. Rest Helps Performance

Several studies have examined the effects of rest in mitigating decrements in performance with time on task (Bergum & Lehr, 1962; Ross, Russell, & Helton, 2014; Helton & Russell, 2015, 2017). For example, Helton and Russell (2015) demonstrated the benefit of rest by having participants carry out a vigilance task (N. H. Mackworth, 1948) that was interrupted by either a rest period or another task before resuming the initial task. In their first experiment, they found that participants who were given a rest period performed better post-interruption than those who remained on task. In a second experiment, Helton and Russell (2015) expanded the set of interruption conditions from two to five (rest, continuation, verbal match to sample, letter detection, or spatial memory). They found that the restorative effects of the interruption was predicted by the degree to which it involved a task that was distinct from (i.e., did not overlap) with the vigilance task: those in the rest condition performed the best post-interruption, and those in the continuation condition performed the worst. Furthermore, participants in the verbal match to sample condition, which had the least amount of overlap with the vigilance task, performed the second best, and participants in either the letter detection or spatial memory conditions (which had partial overlap with the vigilance task) performed better than those in the continuation condition but worse than those in the verbal match to sample condition. Thus, the more that the interruption involved a task similar to the vigilance task, the less it helped.

#### 2. Arbitration Between Labor and Leisure

Kool and Botvinick (2014) proposed that the choice of how much and for how long to engage in a cognitively-demanding task vs. rest reflects a valuation of mental effort and rest as non-substitutable “goods” (i.e., forms of utility), that can be described using the same approach used to analyze the labor/leisure tradeoff in economics (Nicholson & Snyder, 2012). To demonstrate this, they conducted an experiment in which participants were allowed to alternate as they wished between doing three-back and one-back (the latter of which effectively played the role of ‘rest’) versions of the N-back task (Kirchner, 1958) for one hour, with increased time on the three-back resulting in increased compensation. They observed that participants sought a balance between doing the three-back and one-back, which they described in terms of a joint concave utility function combining labor and leisure. Evidence for this concavity came from experiments manipulating the fixed and variable wages, and measuring the direction in which the tradeoff changed. This provided a formally rigorous description of the tradeoff between mental effort and rest, in which fatigue can be interpreted as reflecting the value of rest and, as suggested by the authors, a normative framework for relating the tradeoff to rational models of control allocation (Shenhav et al., 2013) However, this account did not provide an explanation for the value of rest.

#### 3. Difficult Tasks are More Fatiguing

Many fatigue studies that have reported depletion-like effects follow a sequential-task format: Engage the participant in a first task that is ‘depleting,’ and demonstrate there is a negative effect on performing a subsequent second task. For example, Blain, Hollard, and Pessiglione (2016) conducted a study over the span of six hours. Participants performed either the ‘easy’ tasks of a one-back and one-switch^2^ or the ‘hard’ tasks of a three-back and twelve-switch. Every thirty minutes, participants were given a block of intertemporal choice trials. The depletion effect was measured by the amount of discounting in these trials. Although performance on the primary tasks (N-back and N-switch) was comparable across groups and consistent throughout the experiment, participants in the ‘hard’ condition made increasingly more impulsive choices (i.e. discounted more heavily) over the course of the experiment, whereas those in the ‘easy’ condition did not show this effect. Assuming that increased impulsivity reflects fatigue, the results from this experiment suggest that participants in the ‘hard’ condition were more fatigued than those in the ‘easy’ condition.

#### Discussion

Rest thus seems to play a vital role in understanding the normative basis of fatigue. Whereas rest is sometimes assumed to reflect the lack of activity, an extensive body of evidence in the memory literature now suggests that it is a state in which the brain engages in offline processing mechanisms such as planning and consolidation (McClelland, McNaughton, & O’Reilly, 1995; Tambini, Ketz, & Davachi, 2010; Carr, Jadhav, & Frank, 2011; Ólafsdóttir, Bush, & Barry, 2018; Wamsley, 2019). This suggests a grounding for the benefits of wakeful rest — i.e., the value of planning and consolidation, or more particularly the improvement in future reward those processes may achieve. If so, the agent should induce a state of ‘rest’ when its estimated value surpasses the estimated value of physical action^3^. We propose that this value is represented by the phenomenological experience of fatigue. Below, we discuss hippocampal replay as one mechanism of offline processing that has a quantifiable value.

### Hippocampal Replay

Neurons in hippocampus called “place cells” are famously tuned to spatial locations, that is they tend to respond when the organism is in a certain location (O’Keefe & Dostrovsky, 1971; Moser, Kropff, & Moser, 2008). Interestingly, they also fire in coordinated patterns that appear to represent trajectories removed from the animal’s location. Hippocampal replay refers to the physiological phenomenon in which hippocampal place cells fire in sequential patterns during periods of sleep and awake rest (Foster & Wilson, 2006; Diba & Buzsáki, 2007; Davidson, Kloosterman, & Wilson, 2009; Karlsson & Frank, 2009; Gupta, van der Meer, Touretzky, & Redish, 2010). Replay events are commonly observed during epochs of high-frequency oscillatory activity in the hippocampus known as ‘sharp wave ripples,’. When compared with the spatial locations represented by the place cells, the replayed sequential patterns often correspond to spatial trajectories – both experienced and novel – in the animal’s physical environment (M. A. Wilson & McNaughton, 1994; Nádasdy, Hirase, Czurkó, Csicsvari, & Buzsáki, 1999; Louie & Wilson, 2001; A. K. Lee & Wilson, 2002). Though hippocampal replay has been most frequently observed and characterized in rodents, recent studies have also begun to characterize a corresponding phenomenon in humans during periods of rest (Gershman, Markman, & Otto, 2014; Schapiro, McDevitt, Rogers, Mednick, & Norman, 2018; Momennejad, Otto, Daw, & Norman, 2018; Liu, Dolan, Kurth-Nelson, & Behrens, 2019; Wimmer, Liu, Vehar, Behrens, & Dolan, 2019; Schuck & Niv, 2019; Eldar, Lièvre, Dayan, & Dolan, 2020; Liu, Mattar, Behrens, Daw, & Dolan, 2020).

Importantly, sharp wave ripples – and associated replay – occur one trajectory at a time, when an animal is standing still, resting, or asleep. During active locomotion, hippocampus predominantly represents the animal’s current location (or oscillates a bit ahead and behind it, in sync with a distinct mode of theta-band oscillation in the EEG). This is important in the current context, because it means that hippocampal replay events carry an opportunity cost: they are exclusive of active locomotion. It is likely that this reflects contention for a shared resource: the hippocampal representation of location, which can only represent one location at a time, and thus can’t be used simultaneously to represent physical presence at one location but replay of another.

#### Mattar and Daw (2018)

Mattar and Daw (2018) (henceforth referred to as M&D) investigated the utility of hippocampal replay within a reinforcement learning setting. They proposed that replay acts as the physiological instantiation of a step of model-based value computation over that location (Sutton et al., 1998; Daw et al., 2005). Under this model, replay has the potential to affect the agent’s future behavior, and therefore the potential to increase its expected future reward. The place cells activated during replay events are assumed to correspond to the experiential states the agent is simulating.

Replay has been proposed as a mechanism by which the model-based system can be used to accelerate learning relative to traditional model-free algorithms. While the latter, such as Q-learning, have been proven to converge to the optimal policy after sufficient experience (Watkins & Dayan, 1992), this process can be slow in practice because it relies on interactions with the external world. Replay can be thought of as a mechanism by which simulated experience using the model-based system is substituted for physical experience (Sutton, 1991). This is useful, in turn, because experience is actually playing two roles in an algorithm like Q-learning. It is both interacting with the world to gather information about how a task works, e.g. the location of rewards, but also propagating that information along experienced trajectories to work out its consequences for distal actions. The latter function (though not the former) can also be accomplished by mental simulation. For instance, even once you know the rules of chess completely, it takes further computation to elaborate their consequences for the best moves in particular situations. If this simulated experience is faster than physical experience and/or selected through a priority metric (Peng & Williams, 1993; Moore & Atkeson, 1993), the agent can converge to the optimal policy quicker, and thus increase future reward, as opposed to relying exclusively on physical experience.

M&D derived the value of a single replay event, called the Expected Value of Backup (EVB), and ran a set of simulations under the assumption that agents replay the state-action pair (*s_k_, a_k_*) with the highest EVB at the beginning and end of a trial.

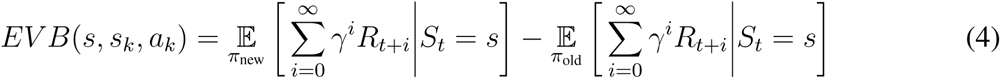

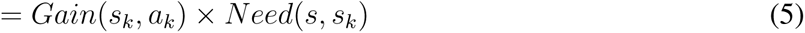

The *Gain* corresponds to the expected increase in expected reward following a visit to the replayed state (since this is the only state in which choice can be affected by a one-step backup) and can be expressed as:

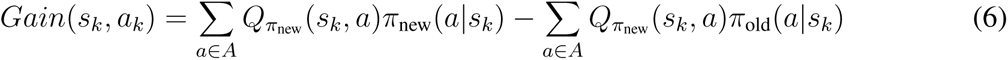

That is, the change in expected future reward *Q* expected following a visit to state *s_k_*, due to following the new policy *π*_new_ resulting from the computation, vs. following the status quo policy *π*_old_. The *Need* term corresponds to the expected number of (delay discounted) future visits to that state:

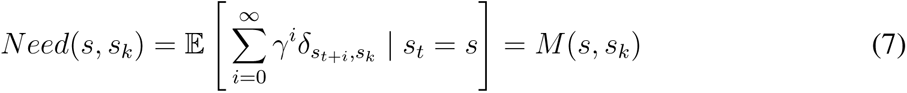

Here, *δ* is the Kronecker delta function, so the Need is the expected future discounted occupancy for the contemplated state *s_k_* starting in the current state *s*. This in turn can be obtained from the successor representation *M* (Dayan, 1993; Gershman, Moore, Todd, Norman, & Sederberg, 2012), estimated as 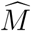, for the state pair (*s, s_k_*).

M&D showed that this model accounts for a wide range of empirical findings in the replay literature, including in particular the reported predominance of forward and reverse replay in the beginning and end of a trial, respectively.

#### Expected Value of Backup with Cost (EVB_*C*_)

While the M&D model provides a rationale for *which* experiences an agent should replay (Peng & Williams, 1993; Moore & Atkeson, 1993; Schaul, Quan, Antonoglou, & Silver, 2015), it does not directly address the question of *when* an agent should replay. Thus, we extend the original M&D model to provide a normative answer this question, by taking into account not only the benefits that replay has for performance, but also the opportunity cost that it carries in time; that is, by delaying the opportunity for reward.

Formally, *EV B_C_* (*s, s_k_, a_k_*) is the expected increase in reward resulting from replaying the state-action pair (*s_k_, a_k_*) while in state *s* and executing the corresponding Bellman backup (i.e. temporal difference learning update, as in Equation 1 but for a simulated rather than experienced step), minus the amount of reward lost due to the time it takes to replay that state-action pair.

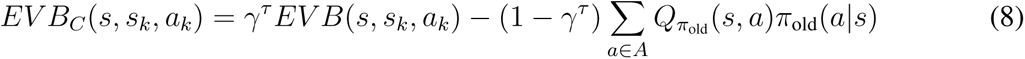

Here, the first term correspond to the Expected Value of Backup (EVB) discounted by *τ* which is the ratio of time it takes to replay versus act^4^. This discounting indicates that the benefits of replay can only be accrued after the time required to replay, *τ*, has elapsed. The second term is the reward lost due to the time it takes to replay: ∑_*a*∈*A*_ *Q_π_*_old_ (*s, a*)*π*_old_(*a|s*) is the expected discounted future reward that would be available if the agent started acting immediately, and *γ^τ^* ∑_*a*∈*A*_ *Q_π_*_old_ (*s, a*)*π*_old_(*a|s*) is the same quantity adjusted by the passage of *τ*. Their subtraction – i.e. (1 *− γ^τ^*) ∑_*a*∈*A*_ *Q_π_*_old_ (*s, a*)*π*_old_(*a|s*) – is thus the reward forgone due to the time it takes to replay. The derivation of this result can be found in the Appendix.

#### Hippocampal Replay as the Value of Leisure

The M&D model, and our subsequent *EV B_C_* extension, provides a quantitative value to ‘rest’ if the agent is engaging in replay during these rest states. Defining *EV B_C_^*^* as max *EV B_C_* (*s, ·, ·*), a rational agent should replay the most valuable location, arg max *EV B_C_* (*s, ·, ·*), as long as its value *EV B_C_^*^ >* 0. Hence, if *EV B_C_^*^* is positive, replaying is more valuable than acting, a situation which we propose is subjectively sensed as fatigue. If *EV B_C_^*^* is negative acting is more valuable than replaying, and thus the agent should physically act instead of resting. Thus, the agent is optimizing the intertemporal tradeoff between acting (providing a more immediate opportunity for reward) and replaying (providing an opportunity for greater but later reward). This insight may help to rationalize the labor and leisure tradeoff that has been described for cognitive control (Kool & Botvinick, 2014; Niyogi, Breton, et al., 2014; Niyogi, Shizgal, & Dayan, 2014; Inzlicht et al., 2014; Dora, van Hooff, Geurts, Kompier, & Bijleveld, 2019).

#### Results

Figure 2 plots the replay behavior of an agent pursuing a specified reward in a gridworld, using the *EV B_C_* -driven replay algorithm descried above, for different values of *τ* (all details of simulations are in the Methods section of the Appendix). Three phases of replay behavior can be seen in these plots. In the first, the agent does not replay because there is no knowledge of the reward structure in the first trial, and thus there is no value to replay. Instead, the agent is accumulating experience to build an internal model of the environment (specifically, it is discovering rewards and developing its successor representation). In the second phase, the agent replays extensively because it has a good internal model but has still not fully developed and refined its value function, and thus can still gain by adjusting its Q-values through replay as well as action. Due to the expanded scope of replay (i.e. the ability to replay *any* experience), the speed benefit of replay, as well the low value of action, *EV B_C_* is positive in this regime. This is the phase we identify with fatigue. Lastly, in the third phase, the agent stops replaying because its value function has become sufficiently good that the opportunity costs of replay exceed its benefits (Van Der Meer & Redish, 2009; van de Ven, Trouche, McNamara, Allen, & Dupret, 2016). This third regime — in which the value function has converged, *EV B_C_* is negative, and behavior is executed without further deliberation — can be thought of as a transition to fully model-free, automatic processing. According to our model, cognitive fatigue, and corresponding periods of rest, arise during the preceding controlled, deliberative phase.

**Figure 2.**
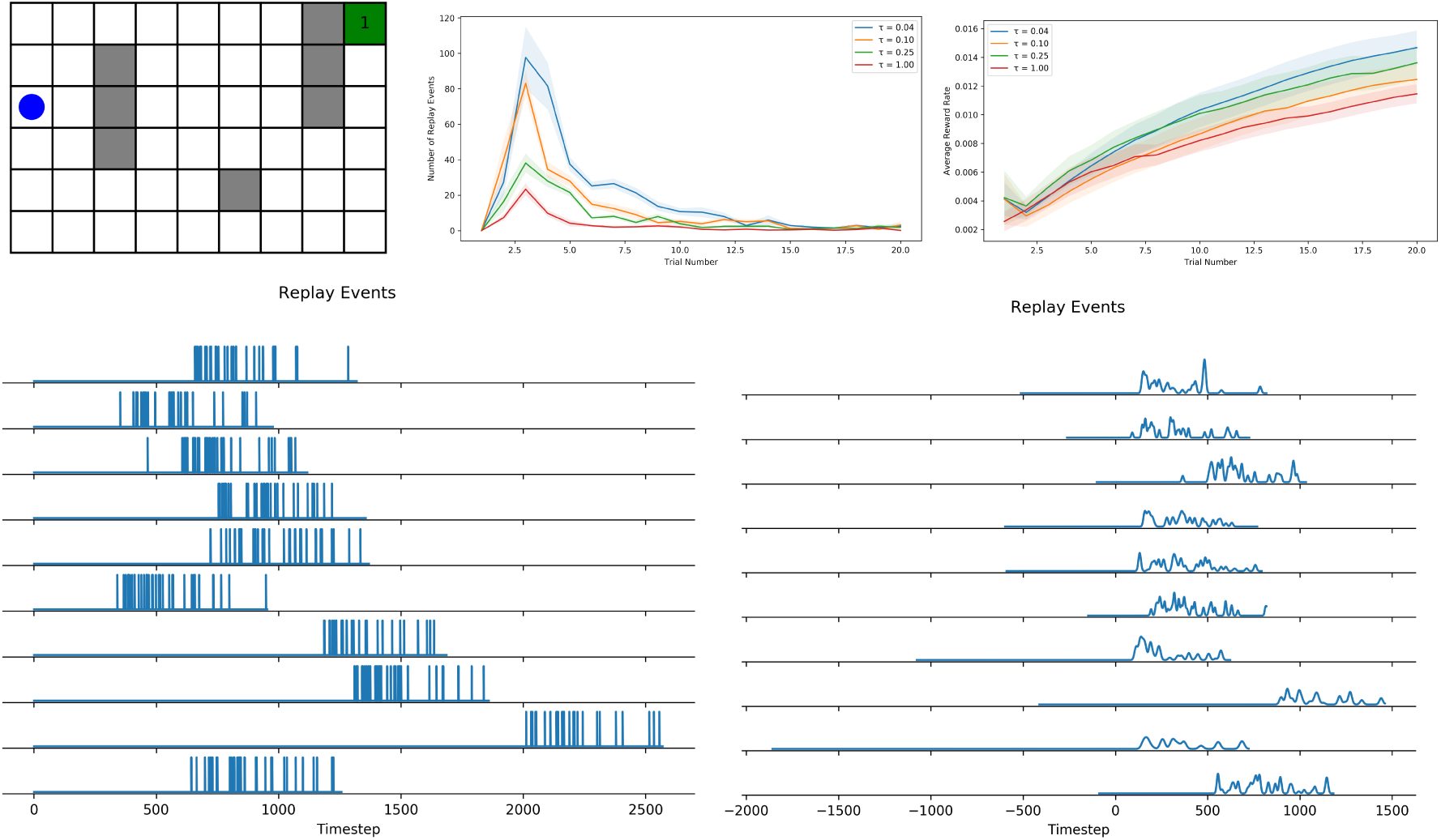
Simulation results for gridworld agent. (Top Left) Gridworld environment used for simulations. (Top Middle) Number of replay events over the course of multiple trials for different values of *τ*. (Top Right) Average reward rate for different values of *τ*. All error bars indicate *±*1 SEM. (Bottom Left) Spikes indicate individual replay events. (Bottom Right) A smoothed version of the left panel. Each trial is also shifted by the amount of time it took for the first trial (which is purely random exploration).

Our model explains the three canonical effects outlined earlier. The restorative power of rest demonstrated in Helton and Russell (2015) can be explained in a straightforward way in terms of replay: rest provided the participants an opportunity for replay that facilitated learning and later performance (and also diminished the need for subsequent replay, which itself would compete with task performance). The second experiment, on the effects of filling a task interruption phase with different interfering tasks, can be explained in the same terms, if it is assumed that the opportunity for replay during the interruption was (inversely) related to the extent to which the task performed during the interruption shared processing resources with those engaged by the vigilance task (e.g., visual encoding and identification of letters). There is strong evidence in the literature that tasks that share processing resources, and risk interference with one another as a consequence, rely on control to mitigate such interference by ensuring that only one is performed at a time (Navon & Gopher, 1979; Meyer & Kieras, 1997; Salvucci & Taatgen, 2008; Musslick et al., 2016). Assuming the same holds for replay (i.e., that it relies on the same perceptual and decision making mechanisms engaged by overt performance), then the more the interruption task shared resources with the vigilance task, the less opportunity it provided for replay of the vigilance task and its salubrious effects.

Figure 2 also grounds the labor-leisure tradeoff of Kool and Botvinick (2014) in normative terms, if it is assumed that leisure corresponds to time allocated for replay. Rather than positing leisure as intrinsically valuable, we demonstrate how the temporal dynamics of its value as an opportunity for replay (mathematically derived in our model) leads it to be either greater than or less than the value of action at different points in time. Accordingly, a rational agent should arbitrate between periods of action and replay, based on which maximizes future reward.

Lastly, to model the relationship between task difficulty and fatigue, we evaluate our agent on an easier gridworld, shown in Figure 3. Since the agent takes less time to develop automaticity in this task (i.e., learn the relevant value function), the overall replay behavior (and hence fatigue) is reduced. Thus, our model is able to demonstrate the relationship between fatigue and task difficulty. Specifically, more difficult tasks are more fatiguing because replay has greater value relative to immediate action in tasks in which fully learning the value function is slower. Therefore, the three-back is fatiguing, as demonstrated in Blain et al. (2016). Conversely, once the value function is learned and the task can be executed automatically without further replay, there is little value in offline processing and thus the model predicts less or no fatigue, as demonstrated in the one-back condition of Blain et al. (2016).

**Figure 3.**
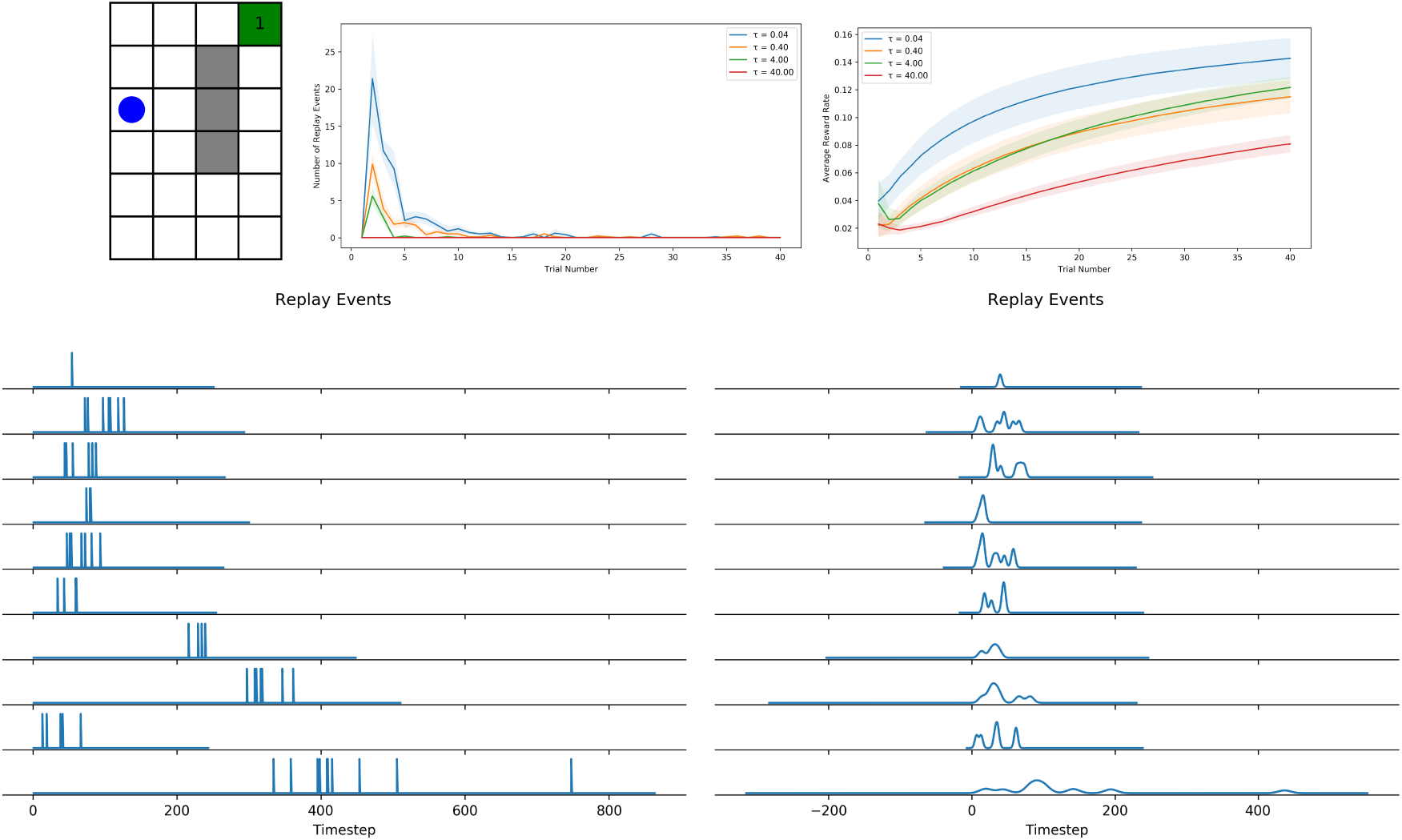
Simulation results for running easy gridworld agent. (Top Left) Easy gridworld environment used for simulations. (Top Middle) Number of replay events over the course of multiple trials for different values of *τ*. (Top Right) Average reward rate for different values of *τ*. All error bars indicate *±*1 SEM. (Bottom Left) Spikes indicate individual replay events. (Bottom Right) A smoothed version of the left panel. Each trial is also shifted by the amount of time it took for the first trial (which is purely random exploration).

### Discussion

The model we propose suggests that the role of leisure goes beyond what it is commonly thought to be “doing nothing.” A large body of evidence suggests that, during states of rest, agents replay past memories to help improve future performance. Rational agents should thus induce these states when they are valued higher than action. We propose that this explains the phenomenological experience, and corresponding behavioral observations, of cognitive fatigue. Furthermore, the need for arbitration between replay and action – as well as the competition between replay and any intervening tasks (such as in the study of Helton and Russell (2015) above) – can be explained as a result of the inability to simultaneously use the same processing resources for different purposes at the same time. Since the purpose of replay is to improve the representations used for action, use of these to replay one set of stimulus-actions sequences while physically engaging in another would produce conflict, and thus both cannot be done concurrently. This is consistent with most hippocampal replay studies to date, which show that sharp wave ripples are rarely observed during locomotion.

#### Benefits and Limitations of Replay

Updating learned action values is one benefit of hippocampal replay, but it is plausible (and probable) that there are other benefits. The complementary learning systems framework (McClelland et al., 1995; Kumaran, Hassabis, & McClelland, 2016; Schapiro, Turk-Browne, Botvinick, & Norman, 2017) suggests that another benefit of offline, hippocampal replay is preventing catastrophic interference that can occur in gradient learning due to the high autocorrelation of online experience (Mnih et al., 2015). Similarly, understanding the extent to which replay during awake rest differs from that during sleep will help inform our understanding of the benefits of the different offline processing mechanisms. There may also be some limitations of replay. Dasgupta, Smith, Schulz, Tenenbaum, and Gershman (2018) proposed that mental simulation may reflect a noisy form of physical simulation. Thus, physical action and experiential learning may be more valuable in situations in which it is difficult to build a model of the environment. However, mental simulation may be more useful (relative to direct trial-and-error learning) for discovering delayed action-outcome relationships in multi-step sequential tasks, such as spatial tasks, social situations, and games. Additional research characterizing different offline processing mechanisms according to these factors will be valuable in generating a more precise understanding of how agents should rationally arbitrate between action and rest states.

#### Intra-Trial Dynamics

Fatigue studies generally consider the number of trials or the time on task as the causally relevant measure. The model proposed here suggests that learning is a mediating variable, and offers a more temporally fine-grained analysis, making quantitative predictions in terms of the states in a Markov decision process. Consistent with experimental observations, some points are better than others for replay within individual trials; replay during rodent navigation tasks most often occurs at the start and end of trials as well as at choice points (Carr et al., 2011; Ólafsdóttir et al., 2018). The dynamics of the *EV B_C_* agent are shown in Figure 4, and thus we predict that sequential tasks should have specific patterns of fatigue dynamics within individual trials.

**Figure 4.**
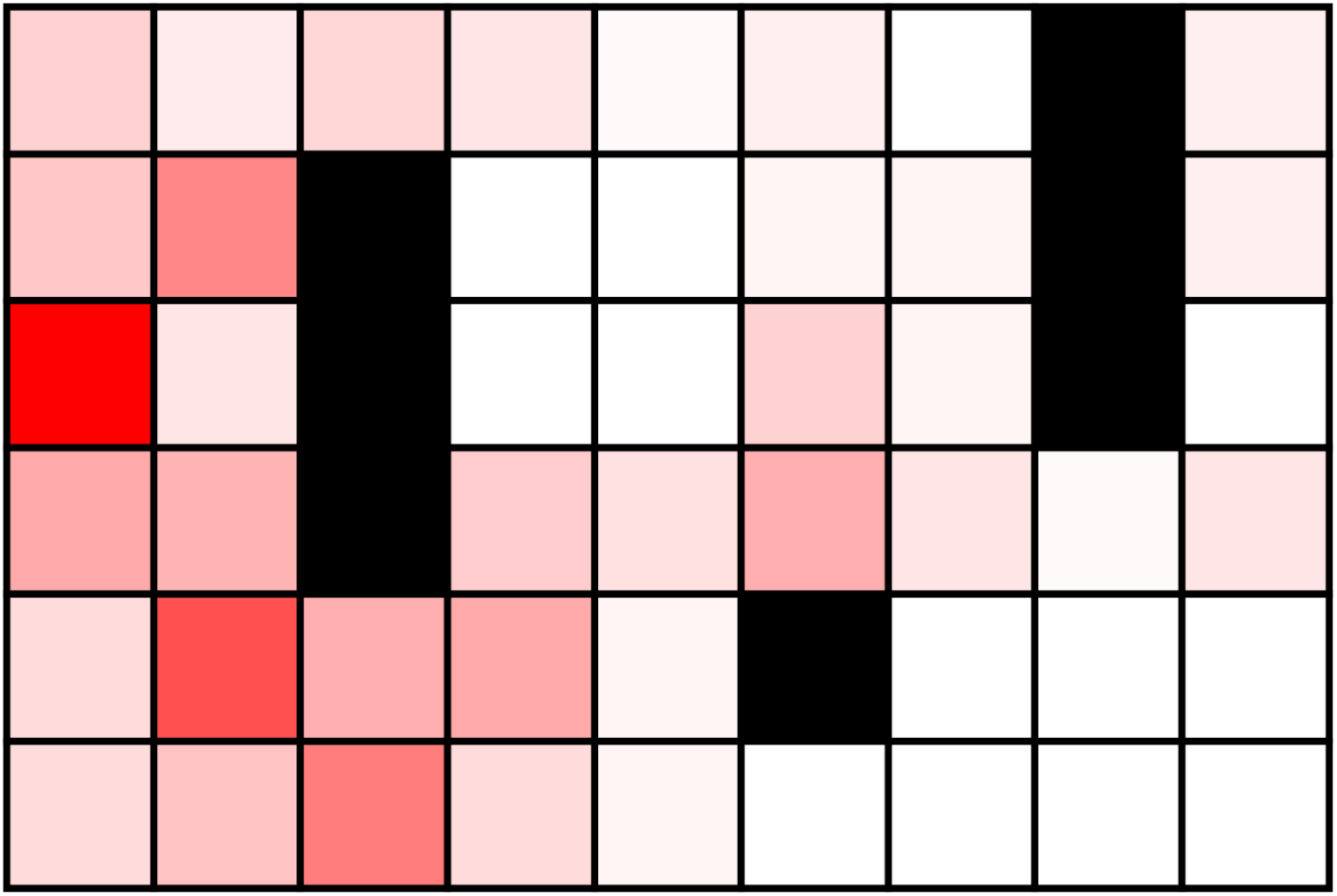
Location of the agent when it decides to replay during a sample run of the original Gridworld environment. Darker red indicates more replay activity. Given that replay is more advantageous in certain states than others, we correspondingly predict that agents will feel more fatigued in some states than others.

#### A Normative Lens to Understand Psychiatric Illnesses

Cognitive control and its attendant costs have been an important focus in the emerging field of computational psychiatry, which aims to give precise mathematical characterizations of mental illnesses in order to increase our understanding of these illnesses and move towards developing effective therapeutics (Montague, Dolan, Friston, & Dayan, 2012; Wang & Krystal, 2014; Huys, Maia, & Frank, 2016). Cognitive control is often considered to be disrupted in many psychiatric illnesses (J. D. Cohen & Servan-Schreiber, 1992; Braver, Barch, & Cohen, 1999), and much of this work has centered around the notion of control costs. Yet, because our understanding of control costs has been so limited, why they are implicated in psychiatric disorders remain unclear. Framing the costs of control as opportunity costs, and specifically by registering the value of replay as an opportunity cost, may therefore be useful in the effort develop a concrete and normative understanding of psychiatric illnesses. Consider post-traumatic stress disorder (PTSD) as an example. If trauma is associated with an event that elicits a particularly large negative prediction error, rational models of memory sampling (Mattar & Daw, 2018; Lieder, Griffiths, & Hsu, 2018) and the current need/gain framework suggest that these should be sampled repeatedly and more often than non-traumatic experiences. This sampling procedure may correspond to the behavioral phenotype of reliving traumatic experiences and rumination. Thus, some symptoms of PTSD may be built on a rational response to negative events, from a rational underlying algorithm being met with an anomalous event (Andrews & Thomson Jr, 2009; Kumaran et al., 2016; Gagne, Dayan, & Bishop, 2018).

#### Hippocampal Lesions

One potential challenge to the *EV B_C_* model would be a hypothetical finding that hippocampal-lesioned patients experience and/or exhibit cognitive fatigue. Although there has not been any systematic study of which we are aware that has measured cognitive fatigue in hippocampal-lesioned patients, it seems likely *prima facie* that this would be observed. To the extent that replay is dependent on the hippocampus, the observation of fatigue in the face of damage to this structure would seem to run counter to the model.

Nevertheless, there are two reasons why this might still be observed. One is the possibility that there are multiple offline processing mechanisms, some of which are hippocampal-dependent but some of which are not, in which case fatigue and the benefits of rest might still be observed even in the absence of the hippocampus. This is quite likely. While we developed our theory referencing hippocampal replay in spatial navigation, which is the case with the most relevant experimental detail, there is a longstanding debate about the extent to which these phenomena are specific to navigation vs. a case of a more general function (G. Cohen & Burke, 1993). Moreover, for other tasks in which deliberative planning has been documented (e.g., multiplayer games and rodent instrumental conditioning) there is at best conflicting evidence of hippocampal dependence (e.g., Corbit, Ostlund, & Balleine, 2002).

The second reason is that, whereas the actual execution of replay may depend (even, for the sake of argument, entirely) on the hippocampus, its engagement is presumably under the control of frontal mechanisms responsible for both monitoring and evaluating the need for replay (Jadhav, Rothschild, Roumis, & Frank, 2016; Shin, Tang, & Jadhav, 2019; McCormick, Barry, Jafarian, Barnes, & Maguire, 2020), and inducing it when needed. This would fall squarely within the scope of theories that suggest frontal structures such as the anterior cingulate and dorsolateral frontal cortex are responsible, respectively, for calculating the expected value of control-dependent processes and engaging those deemed to be most valuable (Shenhav et al., 2013). In that case, whereas a lesion to the hippocampus might impair the ability to *carry out* (and thereby benefit from) replay, it may leave intact the ability to *assess the value* of replay, and the phenomenological correlate of the decision that it is worthwhile (i.e., fatigue). This suggests the intriguing possibility that a double dissociation could be found between distinct contributions of hippocampus and frontal cortex to fatigue.

#### Physical Fatigue

A natural extension of our work is to bring this framework into the domain of physical effort and fatigue. Admittedly, the semantic similarities between cognitive and physical fatigue do not necessarily imply a mechanistic similarity (in fact, some have argued that this perceived mechanistic relationship between physical and mental fatigue have been a distraction; Bartley & Chute, 1947; G. R. J. Hockey, 2011). There are clear physiological components to physical fatigue, which are beyond the scope of the current theory. However, there still may be mental and/or motivational components (Marcora & Staiano, 2010). Whether the effects of these factors are analogous to the role of mental simulation in cognitive tasks may be an exciting direction for future research.

## Part 2: Boredom

The model presented above provides a normative and mechanistic account of the relationship between task difficulty and the dynamics of task engagement, in which fatigue is proposed to signal the value of replay relative to overt task performance. This scales with the difficulty of the task, such that fatigue increases and overt engagement diminishes with greater difficulty. However, diminishing engagement is not restricted to difficult tasks; it is also observed in easy and/or repetitive ones when they are performed for sufficiently long periods of time. This is commonly associated with another phenomenological experience: boredom. That is, after performing even an easy task for enough time, people often experience boredom and prefer to switch to a new task (Bench & Lench, 2013). Here, it seems that overt disengagement reflects disengagement from the task altogether, rather than a switch to a *covert* form of engagement in the service of improving future performance of the current task.

As with fatigue, there is a longstanding literature on the phenomenology of boredom, in which it has been argued that people strive for ‘optimal arousal,’ a state in which stimulation is regulated in order to achieve maximum performance (Yerkes & Dodson, 1908). Optimal arousal theory initially focused on arousal associated with purely environmental stimuli, but subsequent work has suggested that optimal stimulation is also dependent on the individual. For example, ‘flow’ has been described as a state in which an individual is voluntarily and fully immersed in their work (Csikszentmihalyi, 1997). The recently developed MAC model (Westgate & Wilson, 2018) suggests that state boredom (Eastwood, Frischen, Fenske, & Smilek, 2012) is affected by two dissociable components: ‘meaning’ and ‘attention.’ The ‘meaning’ component corresponds to the alignment of the task with the agent’s goals (Van Tilburg & Igou, 2012), while the ‘attention’ component corresponds to an alignment of the agent’s mental resources with the demands of the task (London, Schubert, & Washburn, 1972; Wickens, 1991; Hitchcock, Dember, Warm, Moroney, & See, 1999; Wickens, 2002; Eastwood et al., 2012; Markey, Chin, Vanepps, & Loewenstein, 2014; Raffaelli, Mills, & Christoff, 2018).

Our aim in this Part is twofold. First, we develop a functional understanding of boredom by casting the insights of MAC model in a utility-maximizing framework. Second, we use this formulation to provide insight on the second important issue addressed in this manuscript: the *temporal dynamics* of boredom, specifically *why* boredom seems to increase during easy and/or repetitive tasks. To so, we cast agents in an explore-exploit paradigm, and consider that increases in boredom index the increasing value of exploration (investigating new opportunities that may lead to greater reward in the future) over exploitation (pursuing known, more immediate sources of reward). Mirroring our account of cognitive fatigue, although boredom reflects the relative *value* of other tasks, it is perceived as a *cost* disfavoring status quo action; that is, it indexes the opportunity cost of foregoing exploration by continued engagement in the current task. As we discuss below, the value of information from exploration in the real world (as opposed to the value of mental simulation) captures the remaining puzzles of the relationship between task difficulty and task disengagement.

### Empirical Findings

To motivate our model of boredom, we first summarize the three empirical findings it is meant to explain: (1) boredom is minimized when agents are at an optimal participant-task fit (i.e., the state of “flow”, and the conjunction of meaning and attention), which is often at an intermediate level of difficulty; (2) if agents are bored, they will seek other tasks to perform; and, lastly, (3) boredom increases over time while doing easy and/or repetitive tasks.

#### 1. Optimal Participant-Task Fit

Functional theories consider boredom to arise when the current task is suboptimal for the agent, thus acting as a signal to disengage (Bench & Lench, 2013; Kurzban et al., 2013). Below, we provide a normative interpretation of the MAC model of boredom (Westgate & Wilson, 2018), suggesting that it elucidates situations in which the current task utility is low.

The first component, ‘meaning’, or the relevance of the task for the agent’s goals, directly corresponds to the notion of utility (reward, value) at the heart of reinforcement learning models. Tasks that align with agent’s goals have high utility, whereas tasks that do not have low utility. Low utility tasks such as copying references (Van Tilburg & Igou, 2012), counting words (Geana, Wilson, Daw, & Cohen, 2016a), and passive number viewing (Milyavskaya, Inzlicht, Johnson, & Larson, 2019) are often employed as boredom inductions in the literature. Manipulations increasing the ‘meaning’ of a task, for example by incentivizing performance with charitable donations, reduce boredom even though the task remains the same (Westgate & Wilson, 2018).

The ‘attention’ component corresponds to situations in which there is a mismatch between participant and task: boredom occurs when demands are too high (‘overstimulation’) or too low (‘understimulation’). Tasks that are too difficult have low utility because the probability of success is low (Wickens, 1991, 2002), and thus participants feel bored when required to do a task they cannot do (Fisher, 1987, 1993; Hitchcock et al., 1999; Tanaka & Murayama, 2014). For example, Damrad-Frye and Laird (1989) distracted participants performing a comprehension task with extraneous noise and found that those in the distraction condition felt more bored than those in the no distraction condition.

We propose two reasons why ‘understimulation’ leads to low utility. The first is related to opportunity costs: if an agent is doing an easy task, they can likely perform an additional task, which will have a greater combined utility than doing the sole easy task. Survey results have demonstrated that, to mitigate levels of boredom, many workers perform auxiliary tasks such as reading novels or writing letters (Fisher, 1987). Second, many ‘understimulating’ tasks overlap with those considered to have low ‘meaning’ (e.g. copying references), and thus also have low utility as noted above.

The proposed mapping between ‘meaning’ and utility does not directly address one important question: what are the agent’s goals? That is, while it is generally agreed that copying references, counting words, and passive number viewing are not highly valued tasks, it is not explicitly clear *why* it is the case that an agent’s goals do not align with these tasks. More generally, decision theoretic and reinforcement learning models often view utility as subjective, idiosyncratic to the agent, and do not offer a first-principle account of its source.

A reinforcement learning analysis can offer additional insight, relevant to boredom, as to how other, more distal aspects of a task may contribute to the motivation the agent has to perform the task. In particular, several recent lines of work have started to address this question, by using the value of information framework (Behrens, Woolrich, Walton, & Rushworth, 2007; Bromberg-Martin & Hikosaka, 2009; R. C. Wilson, Shenhav, Straccia, & Cohen, 2019) to suggest that the opportunity for learning is valuable. Agents not only value immediate reward, but also future (discounted) reward, and the value of information quantifies how gaining information, by improving future decisions, can increase expected future rewards when performing a task. Understimulating and/or low ‘meaning’ tasks can often be considered to have a low value of information because there is little to no opportunity for learning available, and, in turn, low value of information for using any such knowledge to attain goals (utility) in the future.

A recent set of experiments by Geana et al. (2016a) directly evaluated this claim. In the first of those experiments, participants were presented with a series of randomly selected numbers from 0 to 100, one at a time, and simply had to predict the next number that would appear. The task was performed in three conditions: in the ‘Gaussian’ condition, numbers were sampled from a Gaussian distribution with a fixed mean and standard deviation; in the ‘Random’ condition, numbers were uniformly sampled between 0 and 100; and in the ‘Certain’ condition, numbers were generated as in the ‘Gaussian’ condition, but participants were told the sampled number before they had to respond, rendering the task trivial. The experiment tested the idea that boredom reflects decreasing information content over time. In that experiment, participants were periodically asked to rate their boredom, and the authors found that this measure was inversely correlated with changes in prediction errors, a proxy for the amount of information being acquired in the task the dynamics of which, in turn, differed between conditions according to what could be learned. A similar relationship between prediction error and boredom has been measured in Antony et al. (2021).

#### 2. Switching to Other Tasks

If the functional role of boredom is to signal the value of disengagement, we should see examples of boredom leading to task switching and general exploration. The second experiment of Geana et al. (2016a) sought to test this directly by allowing participants to switch voluntarily among the tasks used in their first experiment. They reasoned that if boredom is sensitive to the value of information, then it should be possible to demonstrate that participants are willing to forgo reward (i.e., pay) for the opportunity to gain information, by switching to a task that pays less but provides more information. Consistent with this prediction, they found that participants spent the most time in the ‘Gaussian’ condition, in which there was the greatest information content. This behavior runs counter to a standard rational agent model based exclusively on reward, since the ‘Certain’ condition, not the ‘Gaussian’ condition, was the one that maximized current reward. Switching behavior can also be seen in human work environments, in which boredom has been found to lead to a higher labor turnover (Wild & Hill, 1970; Geiwitz, 1966; Kishida, 1973).

Finally, Geana et al. (2016a) conducted a third experiment to test the extent to which opportunity costs associated with task context had an effect on boredom and exploratory behavior. In the first part of the experiment participants performed a standard two-armed bandit task (Berry & Fristedt, 1985) that was used to evaluate their bias toward exploration, during which they also periodically evaluated their boredom. This was followed by an auxiliary task that was known to the participants up front, and was manipulated across individuals to determine the extent to which knowledge of it had an impact on boredom and exploratory behavior in the bandit task. They found that participants anticipating the more interesting auxiliary task reported the bandit task to be more boring and that these self-ratings of boredom correlated with increased exploratory behavior in the bandit task. Interestingly, in this experiment, participants could not voluntarily switch from the bandit to the auxiliary task; thus, taking only that particular situation into account, the value of the auxiliary task should not have had any objective effect on the bandit task. What the results suggest, however, is that boredom may reflect the potential value for exploring alternatives even when these are not immediately or obviously accessible (and, conversely, the opportunity cost of not being able to do so). Taken together, the results of these experiments suggest that the experience of boredom accompanies the propensity to explore, and are consistent with the hypothesis that, more specifically, it signals the estimated expected value of doing so.

#### 3. The Temporal Dynamics of Boredom

Boredom seems to increase over time when performing easy and/or repetitive tasks. Participants in the previously discussed Geana et al. (2016a) study increased their self-report levels of boredom as they engaged in the same task over a number of trials, and a similar effect was measured in Haager et al. (2018). These results support the general idea that tasks should increase in difficulty over time in order to maintain user engagement (Lawrence, 1952; R. C. Wilson et al., 2019), an insight widely leveraged by video games and curriculum designers. Understanding these temporal dynamics will shed insight on arguably the most ubiquitous, everyday experiences of state boredom: we often choose to perform a task precisely because it is not boring, but we eventually become bored of it (and thus choose to switch).

#### Discussion

Boredom can thus be considered as a state in which the current task has suboptimal utility, with the utility function comprised of the defined reward *Q* as well as the value of information *VOI*. A bored agent should then disengage with the present task in order to pursue (or search for) one with higher overall utility, *Q_VOI_* = *Q* + *VOI*. To explain the temporal dynamics, we propose that the change in boredom signals the changing value of information. Once one has mastered a task – that is, one has full knowledge about it, and therefore it has become easy (or as much so as possible) – little remains to be learned that might be of use more generally, making it less valuable to continue and more valuable to move on. This idea has been formalized in models of reinforcement learning (Schmidhuber, 1991; Oudeyer, Kaplan, & Hafner, 2007; R. C. Wilson et al., 2019). In the section that follows, we generalize these formalizations and evaluate the extent to which it contributes to patterns of modulation of performance.

### Formalizing the Value of Information

An approximation to the value of information (VOI) can be expressed generically in a form analogous to EVB in the M&D replay model described Part 1, providing a formal, integrated framework for investigating potential relationships between boredom and fatigue, as follows:

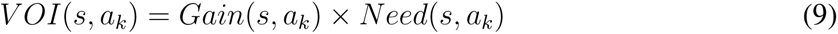

in which

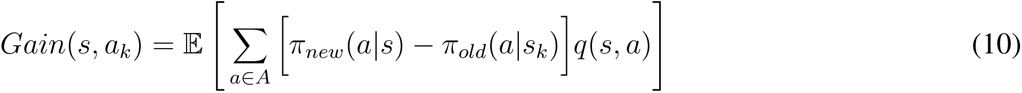

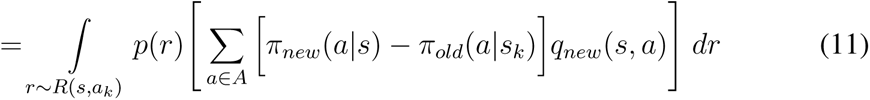

And

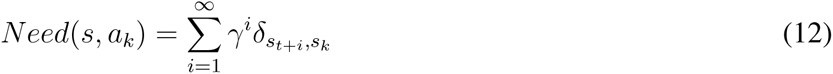

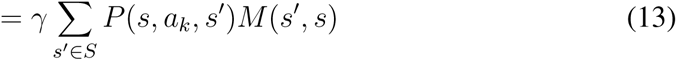

in which *M* is the successor representation (Dayan, 1993; Gershman et al., 2012). The *Gain* term captures the informational value of the obtained reward *r*, as for *EV B* in terms of the resulting change in the choice policy at state *s_k_*. Here this gain will be realized in expectation over which reward is in fact obtained (i.e. over the prevailing prior distribution of *r*). Such gain is obtained following every subsequent visit to *s*, as captured by *Need*.^5^

As an illustrative example, consider a simple two-armed bandit. The first arm starts with a value of zero but, with every iteration, it has a fifty percent chance of increasing by one and a fifty percent chance of decreasing by one. The reward is deterministic based on the arm’s current value. The second arm serves as a baseline and always has a value of one. Here, each arm represents a task, and the stochastic dynamics of the first arm embody a simple form of volatility in the value of options in the world.

Let us assume, as a start, that the agent begins by always choosing the second arm, that is, the stationary option. This allows us to see how the VOI for the alternative, dynamic option changes over prolonged experience with the stationary one. We assume the values for the two actions, *Q_VOI_*, are tracked (as distributions) using simple, recursive Bayesian inference over the true dynamic model of the task.

Figure 5 shows how the action’s *Q_VOI_* values change over time. Notice that, even though the expected one-step reward of both arms remains constant (the first at one and the second at zero), the *Q_VOI_* of the first arm increases over time. As the trials go on, the uncertainty about the first arm’s value increases — as does the VOI for resolving this uncertainty — and choosing it once can give valuable information about which arm to choose in subsequent trials. If the first arm’s actual reward is higher than second arm’s, the agent will change policies after exploring. If it is less, the agent simply goes back to its original policy. At some point, when the uncertainty becomes large enough again (i.e., after choosing the second arm for a sufficient number of trials), the first arm becomes a better choice because of the additional VOI, even though its expected reward (i.e., the mean base *Q* without considering information) is still lower.

**Figure 5.**
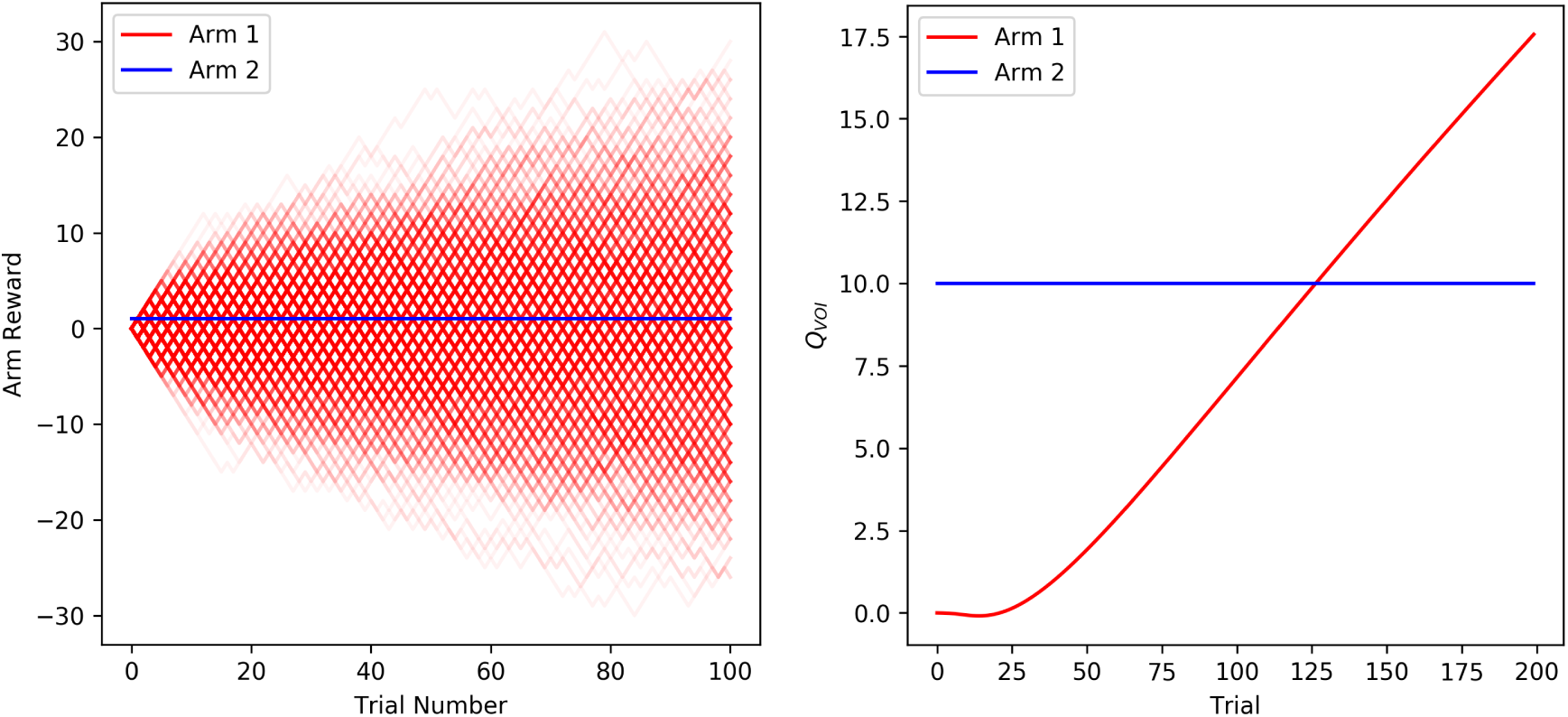
Sample bandit task illustrating the increase in *Q_VOI_* with increased uncertainty. (Left) Red indicates sample reward trajectories of the first arm, the reward for which starts at zero and then increases or decreases by one every iteration. The blue arm 2 indicates an arm with value always at 1. (Right) The change in *Q_VOI_* over time if the agent keeps on picking the blue arm. After a while, the uncertainty of the red arm is large enough that it is advantageous to pick it, even though its expected value is less.

The graph in Figure 5 captures an observation reported in the boredom literature: as one repetitively does a task, the relative value of alternative tasks increases. In our model, this is due to an increase in uncertainty – and therefore the VOI – about the value of the other tasks. This is also coupled with low uncertainty about the status quo task, for which in the current example VOI is always zero because the task is maximally uninformative and static.

The VOI is nonzero when the Q-value posterior is different than the Q-value prior. The increase in value associated with Arm 1 in Figure 5 reflects a situation in which there is learning such that the posterior is different than the prior. In contrast, the value of Arm 2 does not change, since the prior and posterior are never different, and thus there is no gain. This captures the scenario in which a task is too easy (‘understimulating’), as in Geana et al.’s Certain reward condition. The same thing happens for a task that is too hard (‘overstimulating’), e.g. rewards are stochastic but teach you nothing, as in Geana et al.’s Random condition in which there was no information to be gained.

The foregoing simulations describe dynamics of boredom related to uncertainty with respect to its effects on the Gain term. Our formulation also suggests boredom can be affected by the Need term. This can be examined, for instance, by manipulating the horizon in multi-armed bandit settings. For example, Figure 6 plots the probability of choosing Arm 1 in the example above (assuming a softmax decision function), for games of fixed but differing lengths (i.e., numbers of trials), mirroring an effect reported by R. C. Wilson et al. (2014): participants explore more in games with longer horizons, when they have more opportunities to gain information about the uncertain option. In our formulation, this is because the Need term is higher for longer horizons (more expected future choices in which to exploit any learned policy improvements), and thus the overall value of information is higher.

**Figure 6.**
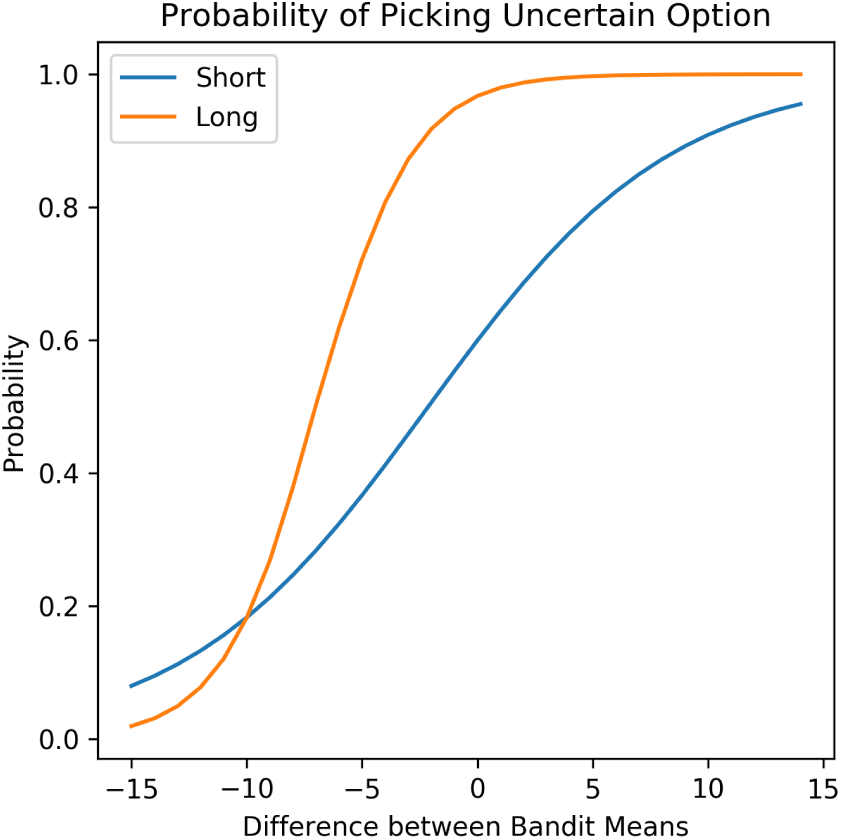
For various mean differences between the two bandit arms, we plot the softmax probability of choosing the option with higher uncertainty in the short horizon versus the long horizon. We reproduce the effect found in R. C. Wilson et al. (2014), in which it was shown that participants in longer horizons are more exploratory. Our formulation provides a parsimonious reason why: the Need term is higher in a longer horizon, and thus the VOI is higher.

### Discussion

We propose that boredom reflects an adaptive signal used to promote exploration and ultimately achieve higher long-run returns. A large amount of evidence has suggested that boredom arises from a suboptimal fit between the participant and the task. Here, we give a formal argument why a suboptimal fit may not maximize long-term reward: when a task permits learning, its true value should reflect additional future gains due to that learning. The optimal task for an individual, then, is one that is rewarding as well as one in which they can continue to learn. Specifically, the agent should take the action with the highest *Q_VOI_*, which balances both the currently known return and the expected increase due to further learning. When these are high, we suggest this corresponds to a *flow* state. This view captures the nonmonotonic relationship between task difficulty and boredom: understimulating and overstimulation (tasks that are boring due to being too easy or too hard) both correspond to situations in which the task’s *Q_VOI_* is low. Furthermore, when the agent is bored, i.e. engaged in a task with a suboptimal *Q_VOI_*, the agent should try to find a task that is a better fit. Below, we discuss different literatures that our formulation can potentially connect to.

#### Adaptive Gain Theory

While our analysis has primarily been at Marr’s (1982) computational level, a complete theory would account for effects at all levels of analysis. One line of work that has pursued both a mechanistic and normative account of arousal has focused on its association with norepinephine (NE) function – a neuromodulator that is widely distributed throughout the brain. The Adaptive Gain Theory (AGT; Aston-Jones & Cohen, 2005) of NE function suggests that low/medium/high levels of arousal correspond to low/medium/high levels of NE release. AGT argues that high levels of NE can be seen as favoring disengagement from the current task set in favor of switching to some other one. Recently, Kane et al. (2017) presented causal evidence supporting this hypothesized relationship. In this work, the authors found that increasing tonic NE levels in locus coeruleus (LC; the brainstem nucleus that is the source of NE) led rodents to explore more in a patch-foraging task. NE levels may thus provide a mechanism by which the brain implements the value of information computation we proposed. Thus, one fruitful line of work would be to link phasic and tonic NE responses to specific exploration algorithms and determine how they change with boredom, in a similar manner to the connection between dopamine and temporal difference learning (Schultz, 1998; Dabney et al., 2020).

#### Curiosity

We have focused on boredom as a negative reflection of the value of information, which drives choices away from uninformative options. It is clearly also the case that there are also affective states associated in a positive way with the informational value of exploration, and may play a complementary role in directing choices toward informative options. Although we have focused on flow, another, that is evidently more directed toward individual alternatives, is curiosity. For instance, Dubey and Griffiths (2019) proposed a model, similar in some ways to ours, interpreting curiosity as an affective state that signals the value of exploring stimuli in terms of their potential for increasing future reward. Thus, one way to dissociate boredom from curiosity is that the former prompts disengagement from uninformative tasks, while the latter prompts engagement to informative tasks.

The emergence of these formal models will hopefully permit future work aimed at determining the extent to which curiosity vs. boredom are simply mirror images, or instead more or less engaged in different circumstances or reflect different aspects of VOI. For instance, we have emphasized the involvement of boredom in driving switching between tasks, although the same informational considerations (and the same models and algorithms) equally apply for within-task learning as well. Are curiosity and boredom engaged at different hierarchical levels of tasks and subtasks, or equally across them? As another example, boredom most obviously relates to the extent of experience with an option, which is a key determinant of VOI. However, a different signal of VOI — novelty — is often invoked in analyses of curiosity. Do boredom and curiosity actually differ in terms of which features of VOI they are sensitive to? Finally, we have argued (and Geana et al. (2016a)’s results support) that boredom is a *differential* signal of *Q_VOI_*, in the sense that it is increased not just by (low) *Q_VOI_* for the current task but also by (high) *Q_VOI_* for alternatives. Is this type of symmetry also true for curiosity?

## Part 3: Replaying to Explore

So far, we have emphasized two different ways in which cognitive or physical actions can be valuable due to their potential for improving one’s future choices. Part 1 modeled how internal computations — replay — can improve learning by propagating knowledge about rewards to distal states and actions. Part 2 modeled the physical actions that gather such knowledge in the first place, and the value of visiting informative (e.g., unexplored) states. We treated these mechanisms as separate and parallel, but they can also interact in important ways. One point of interaction, already reflected implicitly in the dynamics of fatigue in Part 1, is that the value of internal replay is ultimately fed by the gathering of actual information (in the external world) to propagate. However, in many tasks, replay could be used to propagate not only the value of known rewards (e.g., as in our previous simulations, to find the best paths to rewards once they are discovered), but also to propagate the value of information. In fact, one reason exploration in more general sequential tasks (like the gridworlds of Part 1) is more difficult than in bandit tasks (like Part 2) is that in the former, opportunities to obtain VOI can occur at distal states. Reaching those states so as to harvest the VOI itself requires planning, much like figuring out paths to known rewards. Here, we examine the possibility of replay propagating VOI in a combined model, and show that this interaction predicts dynamics of behavior that are different than when replay and exploration are treated separately.

### Simulations

Here, we consider simulations of an agent in a T-maze setting. In this environment, there are two terminal states, each of which has its own reward. Furthermore, the environment is non-stationary, such that every *n* trials the rewards are shuffled between the terminal states. Thus, the agent needs to continuously explore to adapt to the changing reward structure, and must use replay to plan sequential trajectories both to explore and to exploit these rewarding states.

#### Model

We augment the *EV B_C_* model from Part 1 by assuming it propagates gain based on *Q_VOI_* = *Q* + *VOI* rather than reward value *Q* alone. For the current purpose, we also substitute a different approximation for gain in terms of *Q*, based on prioritized sweeping (Peng & Williams, 1993; Moore & Atkeson, 1993):

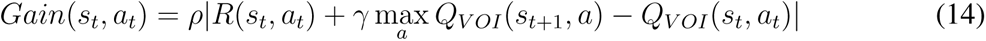

which is the absolute value of the Bellman residual (reward prediction error) at each state. This can be shown to provide an upper bound on the gain as defined previously. This is useful here because, by overestimating gain, it tends to counteract underestimation due to another approximation in our framework. ^6^ We also include an optional degree of freedom *ρ ≤* 1 to scale the heuristic, though we set *ρ* = 1 for our simulations.

Finally, we define a new heuristic for VOI appropriate to the temporal dynamics of reward and resulting uncertainty in these environments (i.e., the rewards shuffle every *n* trials):

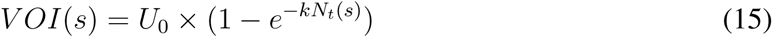

in which *U*_0_ is the maximum value of the uncertainty, *k* is a constant reflecting the hazard rate for switching, and *N_t_*(*s*) is the number of trials since the last visit to the state. *N_t_*(*s*) is initialized as ∞, meaning the VOI is initialized at *U*_0_ and then drops to 0 once visited. After being visited, it exponentially ‘decays’ back to *U*_0_, reflecting the accumulating chance that a change will have occurred.

#### Results

We ran simulations of the agent described above in a T-Maze with different hazard rates (Figure 7). The agent develops uncertainty about the rewarding terminal states, and then uses replay to propagate the corresponding value of information back to the initial states. As a result, we see extensive replay at the beginning, when the agent knows about the existence of (but not yet the value of) the terminal states and propagates its uncertainty through its model of the environment. Then, the agent’s replay behavior follows an oscillatory cycle as the uncertainty about the non-visited terminal state increases and decreases. This exploration is important because rewards are shuffled among the terminal states every five trials. As Figure 7 demonstrates, an agent with this exploration bonus can use replay to achieve a high average reward rate.

**Figure 7.**
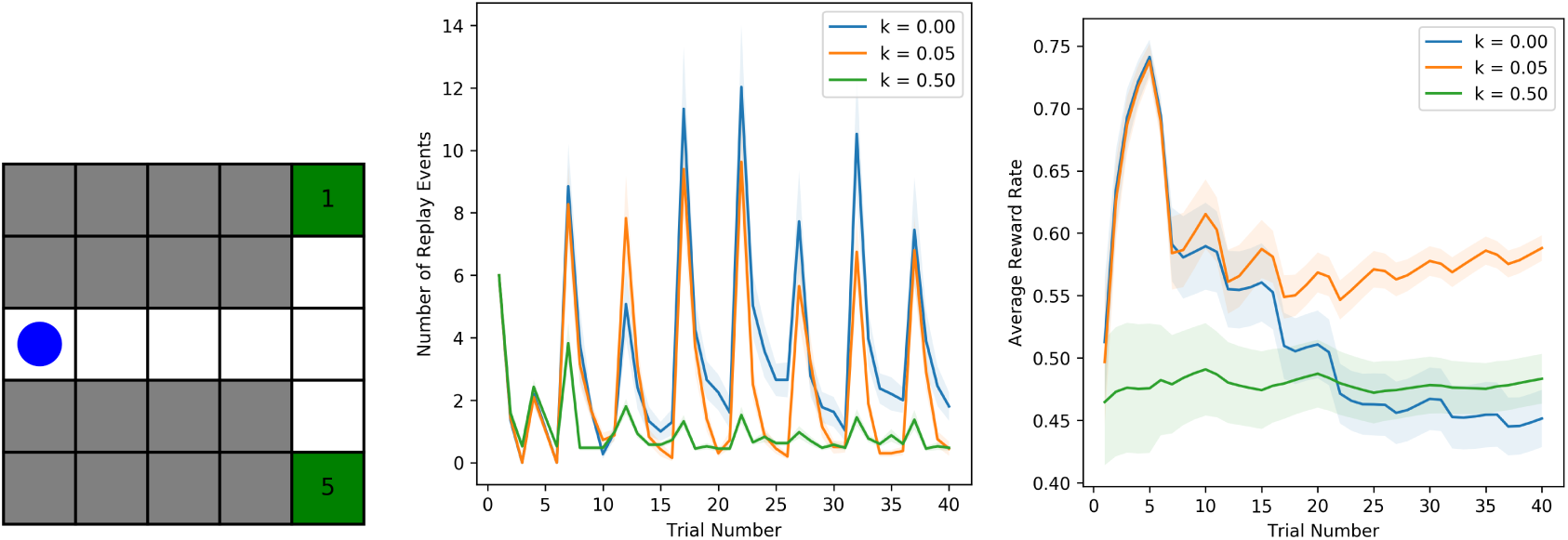
Simulations of an agent that replays and explores. The agent uses replay in order to propagate value of information through the agent’s model of the environment. (Left) Gridworld T-maze environment. (Middle) Replay behavior for different hazard rates. (Right) Average reward rate for different hazard rates. Reward rate is greatest for a moderate hazard rate. All error bars indicate *±*1 SEM.

### Discussion

It has sometimes been questioned whether performance decrements following long time-on-task are a result of fatigue or boredom (J. F. Mackworth, 1968; Pattyn, Neyt, Henderickx, & Soetens, 2008) Meanwhile, other work has not discriminated between these states, implicitly considering both as reflecting motivation and/or opportunity costs (G. R. J. Hockey, 2011; Kurzban et al., 2013). Although the focus of the first two parts of this article was to clarify this distinction, at least hypothetically, by treating them independently of one another, the interaction between replay and exploration considered above shows that there is still ambiguity. In particular, the current simulation predicts two types of replay not present in Part 1: at the beginning of a task, and perpetually even after the task is overtrained. Both of these are due to uncertainty and VOI now driving replay as well as exploratory action. This remains perpetually high because we have assumed (as in the boredom modeling) that the value of terminal states may change if they are long unvisited. Since these relate to both EVB and VOI, it is unclear within our framework which phenomenological state to associate with each.

It is also possible that such interactions could be associated with other phenomenological states. Above, we discussed curiosity as one possibility. Another is ‘mind-wandering,’ the act of spontaneously generating thoughts, that has become a focus of empirical study given its prevalance in daily life (Smallwood & Schooler, 2006; Fox, Nijeboer, Solomonova, Domhoff, & Christoff, 2013; Fox & Christoff, 2014; Smallwood & Schooler, 2015; Christoff, Irving, Fox, Spreng, & Andrews-Hanna, 2016; Danckert & Merrifield, 2018). Recent work has suggested that mind wandering can be goal-directed, and can facilitate creativity (Zedelius & Schooler, 2015; Williams et al., 2018; Fox & Christoff, 2018; Agnoli, Vanucci, Pelagatti, & Corazza, 2018). While no formally explicit account has yet been offered for mind wandering, our framework suggests the possibility that this might correspond to replaying for exploration — a question that invites future research.

## General Discussion

In this article we present a model that provides a formally rigorous, normative interpretation of the phenomena of fatigue and boredom associated with control-demanding tasks. This rests on the widely held assumption that the number of physical and mental tasks that can be performed at once is limited. This imples that engaging in one carries opportunity costs, which our models formalize in terms of the future value of replay for learning (signaled by fatigue) and information gathering through exploration (signaled by boredom). Both of these options involve an intertemporal tradeoff, in that they have the potential to earn greater rewards in the future at the cost of forestalling more immediate reward gathering. This account provides a mechanistic grounding for the intuitive idea that fatigue occurs when performing control-demanding tasks, especially ones that are more difficult (i.e., require more internal computation via replay to learn to perfect); and that boredom occurs when performing easy and repetitive tasks, especially for extended durations (i.e., increasing the likelihood that other, more remunerative opportunities have become available).

The theory predicts that fatigue should track the value of offline processing mechanisms such as hippocampal replay. One of the functions of replay is learning — that is, to facilitate the transfer from controlled to automatic processing — and thus the value of this function is maximized when the agent is still control-dependent. As a result, our account predicts fatigue in control-dependent tasks but not once the agent develops automaticity.

Boredom arises in situations in which the likelihood increases that another more valuable task may be available, favoring the value of exploration. We formalized this exploratory value as the ‘value of information’ and suggested that it tracks the information content available when doing a task. Both easy and impossible tasks offer little information gain, and thus we expect these tasks to be more boring. Boredom is minimized in ‘flow’ states, in which there is both a high exploitative and high exploratory value in the current task.

While this is a subtle distinction, it is a potentially important one, that reflects a primary goal of our effort: to tie the constructs of boredom and fatigue to *distinct* computational mechanisms, in a way that provides a formally precise and empirically testable explanation for two related, but distinguishable ways in which task engagement can decrease over time. This formulation is inspired by, and seeks to explain the subjective phenomenology associated with the terms boredom and fatigue, which we assume reflect computational signals generated by the proposed mechanisms. However, testing this relationship is challenging, given that self-report — the primary means of measuring the phenomenology — may not reliably reflect the underlying mechanisms.

Accordingly, while we hope our theory can explain a sufficient number of findings concerning boredom and fatigue to warrant consideration, it is reasonable to expect that it will not account for all of them. For example, Milyavskaya et al. (2019) surprisingly found that their boredom induction increased participants’ self-report ratings of fatigue more than their effort manipulation did. One possible explanation for this result is that that the self-report scale used ‘fatigued’ and ‘energized’ as the two endpoints, but participants might not consider those to be opposite ends of the same phenomenon. Another tentative explanation is that the bored participants were mind-wandering (e.g., Danckert & Merrifield, 2018), and that the act of mind-wandering increases fatigue. Future work will be required to resolve whether deviations between theory and measures of subjective report in the literature reflect a failure of the theory, or imprecisions in self-report that it may help overcome. Toward that end, one important approach will be to evaluate other forms of measurement that may correlate with boredom and/or fatigue (e.g., pupil diameter or neural signals), and that may provide more proximal markers of the internal computational signals proposed by the theory.

By providing a formal distinction between fatigue and boredom, our model may help guide future empirical work that addresses these phenomena. For example, we previously discussed the multi-hour N-back task from Blain et al. (2016), in which those in the harder task condition exhibited increased delay discounting impulsivity over the course of the experiment. We (concurring with the original authors) interpreted this behavior to reflect fatigue as opposed to boredom. While both conditions are clearly both boring and fatiguing, our computational framework specifies why their relative degree should differ between conditions. When opportunity costs are increasing at a greater rate during a more control-dependent task, fatigue is responsible. If the increase in impulsive choices had instead measured boredom, we would have expected the opposite: those in the easier task condition would change their impulsivity more than those in the harder task. We predict that relatively greater boredom in the easier task condition might indeed be captured using a different dependent measure, such as pupillometry or exploratory choices on a subsequent bandit task. Note also that, under our theory, time discounting would be expected to reflect fatigue only to the extent that patient behavior on delay discounting requires neural operations that compete with replay of the N-back task. While we are not aware of evidence that directly tests this claim, it is consistent with suggestions that intertemporal choice overlaps mechanistically with future-oriented deliberation (Peters & Büchel, 2010; Hunter, Bornstein, & Hartley, 2018), e.g. because patient choices are promoted by adequate mental simulation of their salutary consequences.

Lastly, we conjecture that one reason the line between fatigue and boredom can seem blurred is that replay and exploration can interact. Specifically, agents can use replay to propagate the value of information throughout their model of the environment, thus helping them plan in a way that includes exploration. While this interaction is suggested on purely computational grounds, it raises the possibility that this interaction may be reflected in other forms of phenomenology, such as mind-wandering, offering the potential for a formally rigorous approach to interpreting those phenomena as well.

### Limitations & Future Directions

#### Algorithmic Approximations

Parts 1 and 2 formalized the benefits of replaying and exploring in reinforcement learning environments, but the calculations may not always be feasible. For example, the Mattar and Daw (2018) model (and our version of it in Part 1) computes the value of all replay events before replaying the highest valued event. Such simulations allow us to expose the characteristics of optimal replay, but this is not viable (and not meant) as a realizable process-level account since the selection would take more computation than the computation being prioritized. More realizable, but more approximate, heuristics such as the prioritized sweeping formulation used in Part 3 have been proposed in the computer science (Peng & Williams, 1993; Moore & Atkeson, 1993; Schaul et al., 2015) and neuroscience (Momennejad et al., 2018) literatures, but it remains an open question as to how the brain ultimately computes these values. Similarly, the value of information metric relies on an integral which is intractable in most tasks. Agents may approximate this metric through heuristic algorithms such as UCB (Auer et al., 2002). An alternate approach might be to use learning rate (R. C. Wilson et al., 2019), a heuristic which has improved performance in machine learning environments (Schmidhuber, 1991; Şimşek & Barto, 2006).

#### Causal Disruptions of Fatigue

Although recent empirical studies have begun to test the extent to which boredom plays a causal role in signaling the value of exploration (e.g., Geana et al., 2016a; Geana, Wilson, Daw, & Cohen, 2016b), there has not yet been a direct test of the extent to which cognitive fatigue signals the value of replay. Part of the problem lies in creating an adequate control that would rule out metabolic resource theories. For example, simply stopping participants from taking a break in order to prevent replay would not be an informative manipulation, because a metabolic resource theory would also predict these participants to be fatigued. A more direct test would involve disrupting mechanisms of offline processing in humans (e.g., using transcranial magnetic stimulation, or direct current stimulation), similar to the disruption of sharp wave ripples in rodents (Girardeau, Benchenane, Wiener, Buzsáki, & Zugaro, 2009; Jadhav, Kemere, German, & Frank, 2012), and measuring its impact on performance, reports of fatigue, and inclinations to rest. This represents an important direction for future research.

### Conclusion

The opportunity costs associated with cognitive control exhibit complex temporal dynamics. In this article, we proposed two mechanisms that give rise to these costs: mental simulation (replay) and exploration, that are tracked by fatigue and boredom, respectively. Both reflect an intertemporal choice agents must make in the pursuit of reward maximization. We explained how the independent dynamics of these mechanisms may unify a range of disparate findings in the literature on cognitive control, and proposed that they might interact in a novel way, enabling agents to plan to explore. More generally, they help place cognitive control in the context of approaches, such as bounded rationality and resource rationality (Simon, 1972; Howes, Lewis, & Vera, 2009; Shenhav et al., 2013; Lieder & Griffiths, 2020), that assume agents optimize their utility functions based on a cost-benefit analysis, constrained by the resources and time they have available for computation and action. We hope this provides a useful foundation for future work involving both experimental tests and refinement of theory.

## Acknowledgements

We thank James Antony, Ishita Dasgupta, and Rachit Dubey for helpful comments on previous drafts. This work is supported by the John Templeton Foundation. The opinions expressed in this publication are those of the authors and do not necessarily reflect the views of the John Templeton Foundation. M.A. is supported by the National Defense Science and Engineering Graduate Fellowship Program.

## Appendix

### Methods

#### Part 1: Cognitive Fatigue

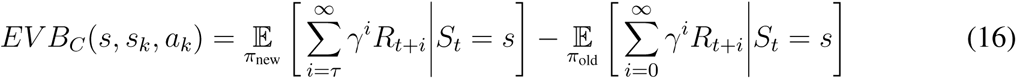

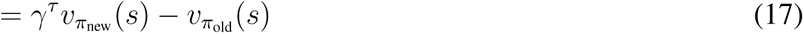

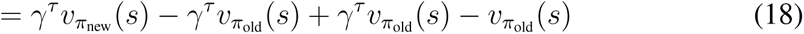

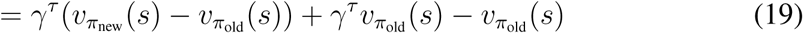

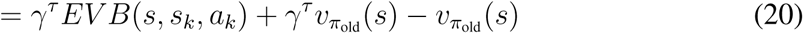

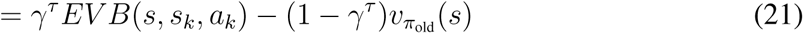

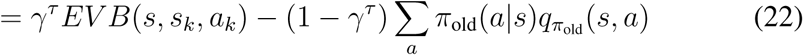

A previous version of the manuscript incorrectly decomposed *EV B_C_* into *Gain × N eed* in which

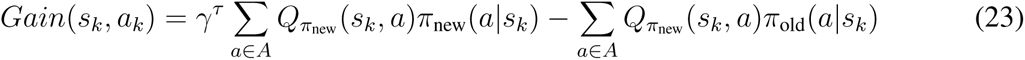

This mistake has been corrected in the present manuscript.

The original gridworld comprised of a 9×6 maze, in which the agent started at (0,3) and there was a reward of 1 at (8,5). Walls were additionally located at (2,2), (2,3), (2,4), (7,3), (7,4), (7,5), and (5,1). The agent’s Q-values and reward prior were both initialized to 0.

An agent’s policy is calculated using the softmax choice rule:

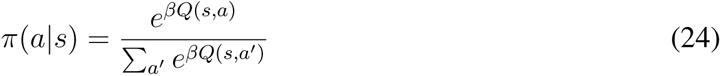

in which *β* is the inverse temperature. *β* was set to 5 for all simulations, unless otherwise stated.

After every state-action-reward-state transition, the agent updates its Q-values according to the temporal difference learning rule in Eq. 1. The learning rate *α*, was set to 0.90 for all simulations. Each replayed backup follow the same update equation. After every actually realized state-action-state transition (i.e., not replayed transitions), the successor representation also updated according the temporal difference learning rule.

At each time step, the *EV B_C_* is computed. If the value is greater than zero, the agent replays arg max *EV B_C_*. Once it falls below zero, the agent samples from its policy *π*.

The easier gridworld comprised of a 4×6 maze, in which the agent started at (0,3) and there was a reward at (3,5). Walls were additionally located at (2,2), (2,3), and (2,4). All other details were the same as the agent in the harder gridworld.

The agent completed twenty episodes for the harder gridworld and forty episodes for the easier gridworld. Ten different runs were completed for each value of *τ*, and number of replay events and average reward rate were averaged over these runs.

The bottom panels in Figures 2 and Figures 3 plot the replay behavior of the ten different agent runs when *τ* = 0.04. The right plot was a smoothed version of the left plot, using a Gaussian filter with *σ* = 5.

#### Part 2: Boredom

The right panel of Figure 5 demonstrates how the *Q_VOI_* of a dynamic task changes over time. An infinite horizon and a discount factor of *γ* = 0.9 were used for simulations. The *q_new_* was calculated using the temporal difference learning update rule.

Figure 6 contrasts the exploratory behavior of an agent in a short horizon versus that in a long horizon. The uncertain action had a relative mean value represented by the x-axis, but was given uncertainty by simulating a fifty percent chance of changing by plus one and fifty percent chance of changing by minus one for twenty iterations. For the short horizon, a horizon of two was used, while a horizon of ten was chosen for the long condition. A discount factor of 0.5 was used for this simulation.

Furthermore, in order to keep the scaling of the Q-values consistent, the policy was computed with respect to the expected one-step reward (which was computed by dividing the Q-value by the Need term). Because of the nature of the bandit task, the Need term for both simulations were calculated analytically instead of using a successor representation.

#### Part 3: Replaying to Explore

The T-Maze was constructed in the Gridworld environment, using a 5×5 grid with walls everywhere except for the middle row and the last column. The agent started at (0,2), and there were two rewards, 1 and 5, at locations (4,0) and (4,4).

Rewards were shuffled randomly every five trials. A successor representation based on a uniform policy *π* was used instead of a dynamically updated successor representation. The agent’s Q-values were initialized to 0, but the VOI’s were initialized to *U*_0_ = 5. For the Need term in Eq. 15, three different values of *k* were used: 0, 0.05, and 0.5. A value of *τ* = 0.04 was used for all simulations. Forty simulations of forty episodes were run for each value of *k*.

## Footnotes

1 While the effects of massing vs. spacing of practice have been studied in several contexts, and several relevant factors have been identified (Izawa, 1971; Bloom & Shuell, 1981; Rea & Modigliani, 1985; Donovan & Radosevich, 1999; Metcalfe & Xu, 2016), fatigue and/or boredom are almost certain to be dominant ones in this example.

2 In an N-switch block, the participant switches between two tasks *N* times.

3 A similar argument has been made for sleep, suggesting that sleep is the ‘price that the brain pays for plasticity’ (Tononi & Cirelli, 2003, 2006, 2014)

4 One estimate of *τ* is 0.04, that comes from the speed of sharp wave ripples in the hippocampus; these occur at approximately 1,000 cm/s (Pfeiffer & Foster, 2013) as compared to the speed of running on a track, which is approximately 40 cm/s (Wikenheiser & Redish, 2015). In humans, a recent study by Wimmer et al. (2019) suggests that replay can be sixty times faster than physical action.

5 These expressions give a partly myopic approximation to VOI: Although they measure the expected future value of exploiting any learning over repeated future visits to *s*, they do not consider the additional informational value of additional learning at these subsequent steps. This approximation was chosen to match the same simplification as used for *EV B* by M&D, as in the previous section, and is sufficient for our purposes.

6 Underestimation arises because *EV B_C_* is defined myopically: it fails to account for the value of a single replay operation in permitting subsequent steps of replay. Accordingly, another way to mitigate this problem would be to extend the original replay model to allow *n*-step backups and calculate their value accordingly.

